# *Blochmannia* endosymbionts reduce brood rearing success in a carpenter ant (*Camponotus* sp.)

**DOI:** 10.1101/2022.12.01.518707

**Authors:** Anika Preuss, Peter Czuppon, Ulrich R. Ernst, Jürgen Gadau

## Abstract

All ants of the species rich genus *Camponotus* (‘carpenter ants’) possess the obligate intracellular bacterial mutualist *Blochmannia*. We tested the relevance of the endosymbiont *Blochmannia* for offspring rearing using cross-fostering experiments between *Camponotus* sp. colonies and subcolonies (worker groups), which were either treated with antibiotics to remove *Blochmannia* or untreated. Our antibiotic treatment reduced the level of *Blochmannia* endosymbionts in eggs, larvae and workers significantly. Corroborating previous results, we found that eggs from treated colonies had a significantly reduced probability to develop into larvae and almost zero probability to become adults. Surprisingly, subcolonies treated with antibiotics had a significantly higher success in raising their own and foreign eggs from treated and untreated colonies than untreated subcolonies. This might indicate that the *Blochmannia* symbiosis entails significant costs for the host in terms of brood rearing, i.e., symbiont-free workers are more successful in brood rearing than untreated workers. If confirmed, this would be a rare case where the costs of a symbiosis can be empirically measured and quantified. Alternatively, the antibiotic treatment increased as a side effect the brood rearing effort of workers leading to the differences in brood rearing success of treated workers. But even if that would be the case, it still indicates that workers that have either lost or have a significantly reduced number of endosymbionts can still raise brood from antibiotic-treated and untreated colonies better than untreated workers. Thus *Blochmannia*, although crucial for brood development, may reduce the amount of brood a colony can raise.

## 1. Introduction

Insects are one of the most successful animal taxa regarding species diversity and geographic distribution. Their evolutionary success might be facilitated by mutualistic relationships with intracellular bacterial endosymbionts (1,2), whereby mutualism is defined as a reciprocal and beneficial interaction between organisms (3). This mutualism ideally results in significant new capacities, which an organism would not be able to realise by itself (4).

The mutualistic relationship with intracellular endosymbionts of some insect species may have granted them access to novel ecological niches because of their tolerance to nutritionally deficient diets (4). This strategy might represent a driving force of insect evolution and a key to the success of these taxa (5). Approximately 20% of all insect species are hosts to intracellular endosymbiotic bacteria, which provide them with essential nutrients that are deficient in the insects’ diet (6). The endosymbionts are tightly interconnected with their host’s metabolic processes, e.g. recycling of nitrogen and provision with vitamins and essential amino acids (7–11). Among Hymenoptera only members of the family Formicidae (ants) are well-known for possessing endosymbionts in bacteriocytes (12–14). These bacteriocytes are specialized cells, which are intercalated between midgut and ovaries and are closely linked to the host’s development (4,15–17).

One of the best-studied mutualistic relationships between endosymbionts and ants occurs among the ant tribe Camponotini (carpenter ants) and the bacterium *Blochmannia. Blochmannia* species are obligate intracellular endosymbionts of ants and were first discovered in the genus *Camponotus* in 1887 by Friedrich Blochmann (18). Preceding phylogenetic studies imply that these endosymbionts were first transferred horizontally from a group of secondary symbionts of mealybugs to the first common ancestor of the Camponotini about 51 million years ago (19–21).

The *Blochmannia* genome contains genes for biosynthesis pathways for essential amino acids (except arginine) necessary for the host’s development, especially for sclerotization, recycling of ammonia, and sulfate reduction (22–24). In contrast, biosynthesis pathways for non-essential amino acids have been lost (20,23–25). This supports the hypothesis that the mutualism between host and *Blochmannia* has a nutritional basis (1,24). In turn, ants provide their endosymbionts with a protected environment within their bacteriocytes allowing a maternal transmission route of *Blochmannia* through the germline (16,26– 28).

General importance of *Blochmannia* for its host has been measured in preceding experiments when the colonies’ success in raising brood has been used as a fitness measure. In contrast to earlier assumptions, treatment with antibiotics did not affect adult workers (29). However, it dramatically impacts larval and pupal development: Ant groups treated with antibiotics had significantly reduced success in raising their own and foreign brood. The brood is fed via trophallaxis and it is assumed that food provided by adult workers is of lower quality due to the absence of endosymbiotic bacteria (29).

Zientz and colleagues (29) analysed the workers’ ability to raise brood, yet they missed to ask whether the lower success in raising brood might not be due to intrinsic factors of the brood itself, e.g., fewer endosymbionts? As previously shown, bacterial transcriptional activity and bacterial genome copy number increases during development, peaks during pupation and young workers, and declines with age in adult ants (29,30) with the greatest increase during embryogenesis and larval development (16,29–32). *Blochmannia* has co-speciated with its hosts and the endosymbionts are transferred vertically (4,15,17,33). Hence, it could be that the observed lower survival of the brood might be due to the brood’s lack of endosymbionts rather than due to lower food quality provided by *Blochmannia*-free adult workers. To answer this question, we conducted cross-fostering experiments and additionally tested if untreated colonies are able to raise brood from treated colonies (29). We found that brood from treated colonies had much lower survival rates, both when raised by workers from treated or untreated colonies, than brood from untreated colonies. Surprisingly, workers from treated colonies were significantly better in raising brood from either treated or untreated colonies than untreated workers, indicating potential costs of the endosymbionts for the host in terms of brood rearing.

## 2. Materials and methods

### Ant culture

Fourteen mated queens of *Camponotus* sp. were collected in the field in April to June 2018 in Comoé National Parc, Ivory Coast (8° 46′ 11″ N, 3° 47′ 21″ W). Within one year, each queen produced a nest with 150-200 workers. Colonies of *C*. sp. were kept in artificial plaster nests in a climate chamber at the University of Münster, Germany, at 25 °C (20-8h) and 29 °C (8-20h), 60% humidity and a 12h day-night rhythm. Twice a week, they were fed *ad libitum* with honey water (50% honey, 50% water), half a cockroach (*Blaptica dubia*) and water. Voucher specimens of workers from our laboratory colonies have been submitted to the Museum für Naturkunde in Berlin.

### Antibiotic treatment

To remove endosymbionts, half of the colonies (i.e., 7 out of 14 colonies) were fed twice a week for 12 weeks with the antibiotic rifampicin (Serva Elektrophoresis GmbH, Heidelberg, Germany), dissolved in honey water (50/50) at a final concentration of 1% (10 mg/ml).

### DNA extraction and quantitative PCR

We used qPCR of bacterial DNA to quantify the *Blochmannia* levels in brood and workers. For each of the seven nests treated with antibiotics, we collected 10 eggs (pooled sample) and two workers before and after the treatment, for a total of 28 worker samples and 14 samples of 10 pooled eggs. We extracted DNA using the QIAGEN QUIAmp DNA Mini Kit (QIAGEN, Hilden, Germany), following the manufacturer’s instructions (QIAGEN 2018) and quantified the amount of *Blochmannia* DNA using 16S rDNA primers (F 5’-AAACCCTGATGCAGCTATACCGTGTGTG-3’, R 5’-CCATTGTAGCACGTTTGTAGCCCTACTCA-3’). As a control, we used 18S rDNA (F 5’-AGGCAGTTAARGAAATTCAA-3’, R 5’-TATTGTCCAGWCAYTACGGGARKC-3’) and 28S rDNA (F 5’-AAGCTAAVCAGAAAGCGGGGA-3’, R 5’-AAAACCATTCGTCTTGACCRC-3’) primers to the host’s rDNA. QPCRs were performed using a KAPA SYBR FAST kit (Merck, Darmstadt, Germany) according to the manufacturer’s instructions. In each qPCR, 0.5 μl DNA template was used. Quantification was performed based on independent DNA preparations and was measured in duplicate.

### PCR *Wolbachia*

We used *Wolbachia* specific primers (16S: F 5’-CATACCTATTCGAAGGGATAG-3’, R 5’-AGCTTCGAGTGAAACCAATTC-3’ (34); wsp: Forward 5’-TGGTCCAATAAGTGATGAAGAAAC-3’, Reverse 5’-AAAAATTAAACGCTACTCCA-3’ (35) to test for the presence of *Wolbachia* (protocol see (36)).

### Cross-fostering approaches

Colonies treated with antibiotics produce fewer pupae than untreated colonies (29). This might be due to (i) less viable brood, and/or (ii) lower ability of the nurses to raise brood. To test hypotheses (i) and (ii), we set up a full-factorial cross-fostering experiment.

From each of the 14 colonies (7 treated with antibiotics, 7 untreated) four queenless subcolonies containing 20 workers were separated, resulting in a total number of 28 subcolonies from untreated (“Untreated”) and 28 subcolonies from antibiotic-treated mother colonies (“Treated”). Within 4 weeks, each subcolony received two times 20 eggs: once eggs from an antibiotic-treated colony (“Treated”), once from an untreated colony (“Untreated”) (Table 1). To test whether workers would discriminate or prefer eggs of foreign colonies, we also compared the acceptance of eggs from the own mother colony (“Same”) with eggs from different mothers (“Different”).

**Table 1:**
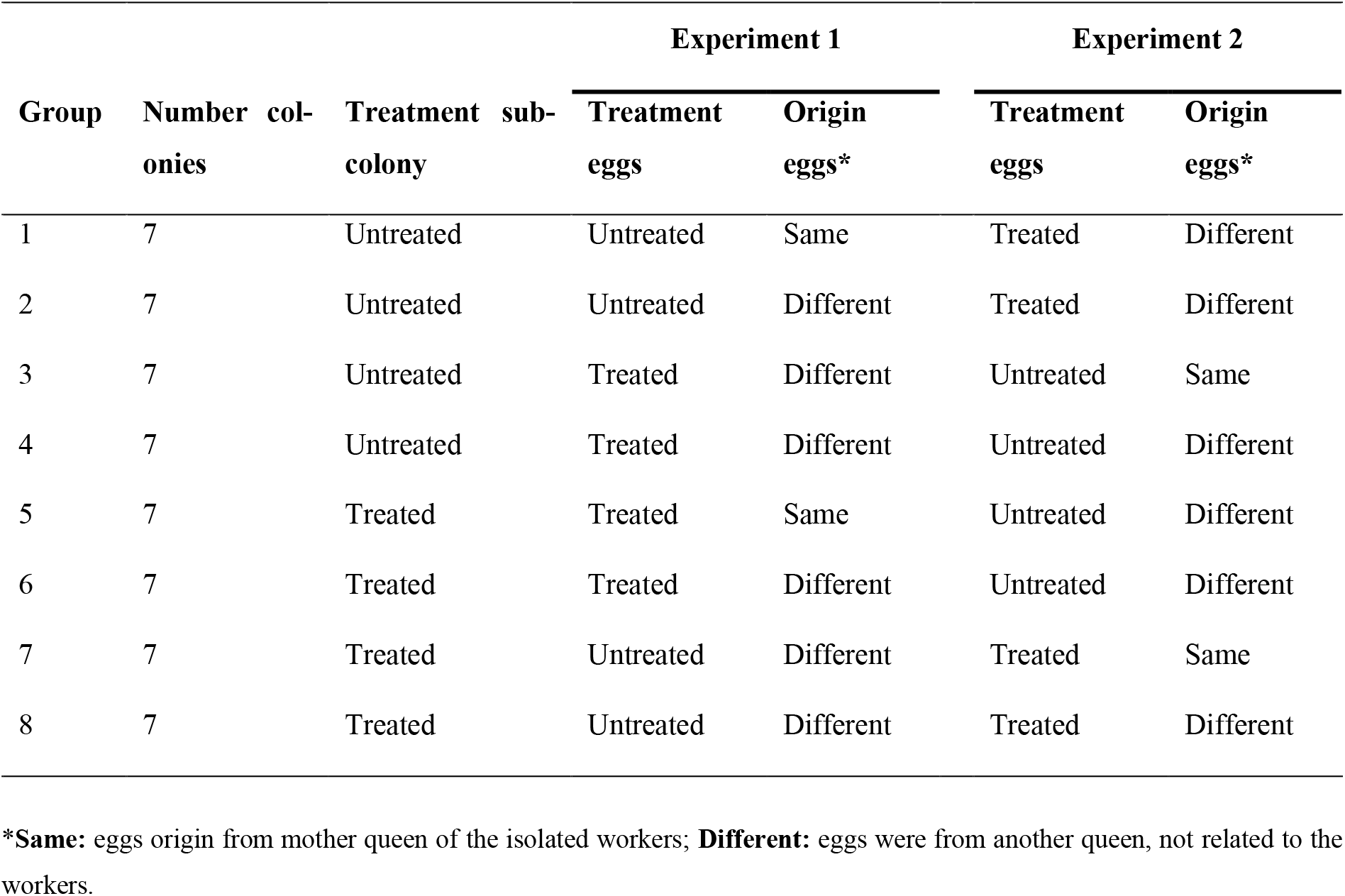
Overview of the cross-fostering approach.

Every day, the number of eggs, larvae, and pupae were counted for each subcolony and pupae were removed from colonies. After 28 days the first cross-fostering experiment (experiment 1) was terminated, and all brood was removed from the subcolonies. The next day we repeated the experiment (experiment 2) and introduced 20 eggs from another colony and treatment.

### Statistical analysis

All statistical analyses were performed using R version 4.0.3 (37). We used Mann-Whitney-U-tests to analyse statistical differences in 16S DNA before and after antibiotic treatment detected by qPCR. For analyses of differences between no treatments and antibiotic treatments, we used generalized linear mixed models with a logit link function (lme4 package; (38)), including random factors for replicates and the origin of ants and eggs used for cross-fostering experiments. We additionally used a penalized maximum likelihood estimation (blme package; (39)) to account for potential problems of data separation (eggs to larvae survival) or overfitting (larvae to pupae survival), details are provided in the comments of the R-script. To assess the quality of the model estimation, we used the DHARMA package (40). The R-scripts to run the analyses are attached as supplemental files.

## 3. Results

We could not detect any signs of a *Wolbachia* co-infection in adult workers. By contrast, all adults and pooled egg samples were positive for *Blochmannia*. Treatment with antibiotics significantly reduced the amount of *Blochmannia* by about 67% in workers (Mann-Whitney-U-test, p = 0.03788), and 99.98% in eggs (Mann-Whiney-U-test, p = 0.0005828) (Supplemental Fig. S1 & S2). In general, eggs from colonies treated with antibiotics were less viable: out of 20 eggs from treated colonies, on average only 6.4 (SD = 5.01, n = 56) brood items (larvae, pupae) were recovered after the end of the experiments (4 weeks). In contrast, 19.8 (SD = 0.93, n = 56) brood items were recovered from eggs that came from untreated colonies. There was a strong and significant effect on survival depending on the ants attending the brood. At the end of both experiments, eggs (n=20) from treated colonies, attended by treated workers, produced 9.4 (SD = 5.25, n = 28) brood items (= 47%), whereas only 3.3 (SD = 2.04, n = 28) brood items (= 16.5%) survived when tended by untreated workers (Figure 1, Table 1).

**Fig. 1:**
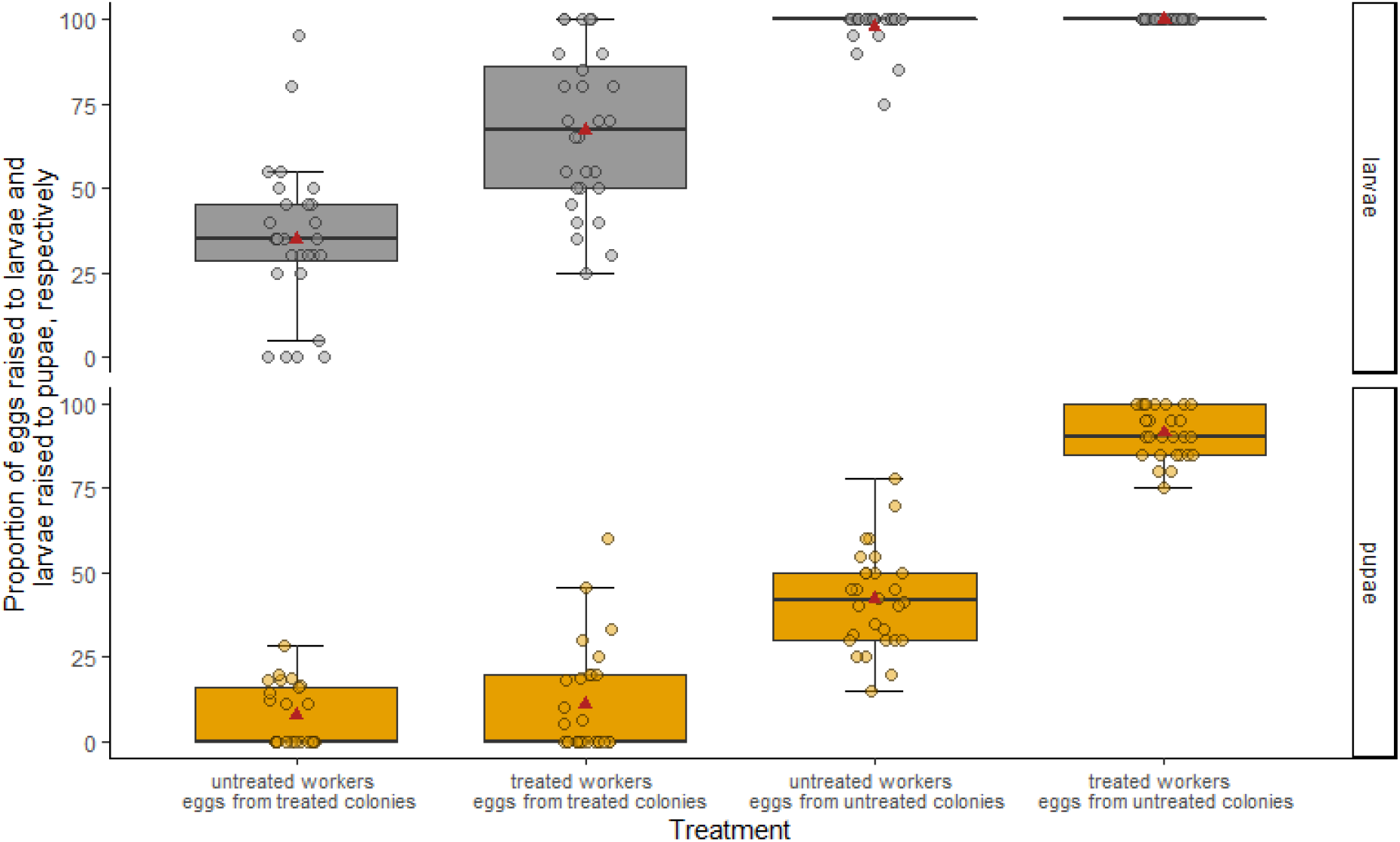
Success rate of rearing eggs to larvae and pupae. Eggs from treated colonies develop less often into larvae and even less often into pupae. Treated worker ants are better in rearing eggs (from treated and untreated colonies) than untreated worker ants. 20 eggs from treated and untreated colonies were taken care by 20 (un)treated workers (full factorial design). Each group of ants raised once eggs from untreated and once from treated colonies. Treatment consisted of antibiotic supplement [rifampicin] in honey water for 90 days. Boxplots show median (thick solid lines), mean (triangles), interquartiles (box boundaries), and 1.5 times the interquartile range (whiskers).

For brood from untreated colonies, we did not find any difference in survival depending on the ants attending the brood: ants treated with antibiotics kept all brood from untreated colonies alive (n = 28); whereas untreated ants kept on average 19.6 items alive (5 out of 28 groups did not manage to keep all brood alive).

When raised by antibiotic-treated workers, 20.0 out of 20 (100%) (SD = 0.0) eggs from untreated colonies developed into larval stage, while on average 13.4 of 20 (67%) (SD = 4.68) eggs from antibiotic-treated colonies could be reared by those workers. In contrast, when raised by untreated workers 19.6 of 20 (98%) (SD = 1.14) eggs from untreated colonies and 7.0 of 20 (35%) (SD = 4.45) eggs from antibiotic-treated queens became larvae.

During the experiments, eggs from antibiotic-treated queens developed slower from eggs to pupae or even not at all in comparison to eggs from untreated queens: In total, 1.3 (6.5%) (SD = 1.76) eggs from treated colonies and 18.3 out of 20 (91.5%) (SD = 1.46) eggs from untreated colonies reached pupal stage when tended by antibiotic-treated workers, whereas only 0.7 out of 20 (3.5%) (SD = 0.98) eggs from treated colonies and 8.3 (41.5%) (SD = 2.91) eggs from untreated colonies were reared by untreated ants. Thus, antibiotic-treated ants were at least twice as successful as untreated ants in raising eggs from antibiotic-treated queens to pupae.

Workers from antibiotic-treated colonies were better at raising eggs to pupae than workers from untreated colonies. Taking the number of hatched larvae as a baseline, instead of the number of eggs at the beginning of the experiment, we observed that 11.1% of larvae deriving from eggs from treated colonies and 91.4% larvae from eggs from untreated colonies could be reared to pupal stage by antibiotic-treated workers. In contrast, only 6.6% larvae from antibiotic-treated queens and 42.2% of larvae from eggs deriving from untreated colonies reached pupal stage when raised by untreated workers.

We detected statistically significant effects of the treatment of eggs and ants on the survival of eggs to larvae (p<2×10^−16^ and p=0.00178, respectively). Untreated ants reared eggs less successfully than treated ants (reduction of 55% in the survival probability from eggs to larvae), and eggs from untreated colonies had a 40% higher survival probability to survive to the larval stage than eggs deriving from treated colonies.

For the survival from larvae to pupae, we found that the generalized linear mixed model with the treatment of eggs and ants as fixed effects did not match the observation well (diagnostic Fig. S1). We therefore included an interaction between the treatment of ants and treatment of eggs, which explained the data better (AIC without interaction = 448, AIC with interaction = 399). This statistical model showed that the treatment of eggs significantly affects the survival from larvae to pupae (p<2×10^−16^). The simple effect of treatment of ants was not statistically significant (p=0.836), but the interaction between the treatment of eggs and ants was (p= 9×10^−15^). Larvae that emerged from eggs from untreated colonies had a strongly increased survival to the pupae stage compared to eggs from treated colonies (more than 700% increase). However, eggs from untreated colonies that were reared by untreated ants suffered from a 50% decrease in survivability from larval to pupae stage compared to eggs from untreated colonies that were reared by treated ants.

We included random effects for the replicate number, the origin of the eggs and ants used for the experiment, to account for potential variation between the colonies (origin eggs and ants) and potential temporal variation during the experiment (replicate). Interestingly, we found that the origin of the eggs and ants had a much stronger effect on the survival probability from eggs to larvae than from larvae to pupae, as visible by comparing the respective cumulative variances, i.e., the sum of the estimated variances for egg and ant origin: egg to larvae = 1.37, larvae to pupae = 0.05. The variation introduced by the replicate remained similar, at least on the variance scale: egg to larvae = 0.18, larvae to pupae = 0.43.

## 4. Discussion

The aim of this study was to investigate the role of the endosymbiotic bacteria *Blochmannia* sp. for brood development in *Camponotus sp*. carpenter ants using a cross-fostering experiment with ant nurses and eggs of untreated and antibiotic-treated colonies. Antibiotic treatment reduced the endosymbiont levels in workers and eggs significantly, which was confirmed by qPCR.

The cross-fostering experiments confirmed that *Blochmannia* is relevant for brood development, since brood with reduced *Blochmannia* titers had a higher egg mortality and slower development rate than brood containing endosymbionts (e.g. (1,16,30)) In particular, these endosymbionts selectively regulate germline genes in early development stages of their hosts to successfully integrate *Blochmannia* into the Camponotini ants and facilitate the horizontal transfer and biosynthesis pathways (19,24,32). By contrast, the frequency of *Blochmannia* decreases in older workers and the endosymbionts seem to play a minor role for adult ants (30).

Surprisingly, we found that antibiotic-treated workers are more successful in raising larvae to pupae than untreated workers, regardless of the origin of the brood. In contrast, there was no effect of the ants on the hatching success of the eggs. During the egg stage, workers provide merely hygienic services and maintain favourable climate conditions but do not exchange information with the egg. However, our statistical model shows a significant interaction between ant origin and the pupation rate. This makes sense given that workers exchange food and information with larvae via trophallaxis.

This suggests that the symbiosis between *Blochmannia* and Componotini ants is more complex than previously thought, revealing a trade-off between benefits (i) and costs (ii) for endosymbionts and hosts at different levels in this symbiotic relationship:

i. *Blochmannia* provides access to nutrients necessary for host development (22–24), while the ants bacteriocytes provide their endosymbionts with a protected environment (16,26–28).
ii. Workers still harbouring *Blochmannia* are less successful in raising brood, possibly due to negative effects on body conditions (resource drainage) in adult life stages or by increasing the susceptibility to pathogen infections and affecting the host’s immune response (41).

It is assumed that the ant’s immune system aims to actively downregulate the number of *Blochmannia* after moulting, revealing a coevolutionary process (arms race or trade-offs) between host and endosymbiont (41,42). By downregulating *Blochmannia* in their bacteriocytes, adult workers are more successful in raising brood. This benefits both, the host and the endosymbiont, since *Blochmannia* is transmitted maternally (and workers do not reproduce), and thus ultimately can only spread if its host reproduces. *Camponotus* sp. ants reproduce via the production of winged queens and males that leave a colony to mate and found a new colony independently. *Blochmannia* endosymbionts are only transmitted via the queens, and the larger the workforce of a colony the more queens it can produce. Therefore, it would be in the endosymbionts very own interest to reduce its prevalence in adult worker ants. That it is not reduced completely may be because the timing and/or mechanism to reduce *Blochmannia* is hard to evolve. Another possibility is that adult ants still benefit from low levels of *Blochmannia* due to their ability to recycle nitrogen from urea (1,23).

We therefore consider the *Camponotus*/*Blochmannia* system as a dynamic symbiosis between endosymbionts and hosts that is more complex than previously appreciated. Further analyses of brood care, immune responses at different development stages, and selection pressures are necessary to improve our understanding of this complex evolutionary system between *Blochmannia* and *Camponotus* ants.

## Acknowledgements

We are grateful to Kathrin Brüggemann and Hilde Schwitte for their help with all laboratory activities, and Jana Salich and Marius Pohl for their support in feeding the animals. Thanks to the entire “Institute for Evolution and Biodiversity”, especially to the “Molecular Evolution and Sociobiology Group”. We thank the Comoé National Park Research Station and its staff for the use of their facilities for field and laboratory research and the park management of Office Ivoirien des Parcs et Réserves for enabling field research in the park. The research was carried out under research permit number N°018/MINEDD/OIPR/DZ from the Office Ivoirien des Parcs et Réserves and adhered to the requirements of the relevant re-search guidelines. The experiments detailed here comply with the current laws of the country in which they were performed.

